# Unbiased K-mer Analysis Reveals Changes in Copy Number of Highly Repetitive Sequences During Maize Domestication and Improvement

**DOI:** 10.1101/078790

**Authors:** Sanzhen Liu, Jun Zheng, Pierre Migeon, Jie Ren, Ying Hu, Cheng He, Hongjun Liu, Junjie Fu, Frank F. White, Christopher Toomajian, Guoying Wang

**Affiliations:** Department of Plant Pathology, Kansas State University, Manhattan, KS. 66506. U.S.A; Institute of Crop Science, Chinese Academy of Agricultural Sciences, Beijing 100081, P.R. China; State Key Laboratory of Crop Biology, Shandong Key Laboratory of Crop Biology, Taian 271018, P.R. China; College of Life Sciences, Shandong Agricultural University, Taian 271018, P.R. China; Department of Plant Pathology, University of Florida, Gainesville, FL. 32611. U.S.A.

**Keywords:** comparative genomics, repetitive, k-mer, maize

## Abstract

The major component of complex genomes is repetitive elements, which remain recalcitrant to characterization. Using maize as a model system, we analyzed whole genome shotgun (WGS) sequences for the two maize inbred lines B73 and Mo17 using k-mer analysis to quantify the differences between the two genomes. Significant differences were identified in highly repetitive sequences, including centromere repeats, 45S ribosomal DNA (rDNA), knob, and telomere repeats. Previously unknown genotype specific 45S rDNA sequences were discovered. The B73-specific 45S rDNA is not only located on the nucleolus organizer region (NOR) on chromosome 6 but also dispersed on all the chromosomes in B73, indicating the relatively recent spread of 45S rDNA from the NOR. The B73 and Mo17 polymorphic k-mers were used to examine allele-specific expression of 45S rDNA. Although Mo17 contains higher copy number than B73, equivalent levels of overall 45S rDNA expression indicates that dosage compensation operates for the 45S rDNA in the hybrids. Using WGS sequences of B73xMo17 double haploids (DHs), genomic locations showing differential repetitive contents were genetically mapped. Analysis of WGS sequences of HapMap2 lines, including maize wild progenitor teosintes, landraces, and improved lines, decreases and increases in abundance of additional sets of k-mers associated with centromere repeats, 45S rDNA, knob, and retrotransposon sequences were found between teosinte and maize lines, revealing global evolutionary trends of genomic repeats during maize domestication and improvement.

## Introduction

The maize genome (*Zea mays* ssp. *mays*) exhibits high levels of genetic diversity among different lines [1–3]. The inbred lines B73 and Mo17 represent two of the most appreciated models for understanding maize genome diversity with respect to small-scale polymorphisms [4–6] and large-scale structural variation [7, 8]. In addition, mapping populations of inter-mated B73xMo17 recombinant inbred lines and double haploids have been generated to facilitate genetic analyses [9, 10]. Numerous comparative genomics studies of other maize cultivars and wild ancestors have examined the origin of maize as well as events of adaptation and artificial selection [11–17]. However, the studies are limited to comparisons of non-repetitive and low-repetitive sequences.

Cytogenetics, genetics, and a few genomics studies have documented variation for many of high repetitive sequences among maize lines, which may also contribute to maize evolution and domestication [2, 18–20]. In maize, highly repetitive sequences are comprised of several major classes, including ribosome DNA (rDNA), knob repeats, centromere satellite C DNAs (CentC), telomere repeats, and various retrotransposon families. The rDNA repeats consist of two classes, 45S rDNA and 5S rDNA, which are transcribed to ribosomal RNAs (rRNAs). 45S rRNA is further processed into 18S, 5.8S and 26S mature rRNAs, which, are then assembled with the 5S rRNA into ribosome subunits [21]. 5S rDNA loci are physically located at the distal of the long arm of chromosome 2 [22], while 45S rDNA tandem arrays are clustered at the nucleolus organizer region (NOR) located at the short arm of chromosome 6 in maize [23]. The copy number of 45S rDNA repeats is highly variable between different maize lines, possibly due to unequal crossover within large tandem repeats [24]. 5S rDNA loci, in contrast, appear relatively stable [25]. Knob repeats are composed of highly condensed heterochromatic regions and are cytologically visible on chromosomes of maize and the wild relatives. Knobs consist primarily of a 180 bp repeat as well as a second 350 bp repeat, the TR-1 repeat. Both types of repeats are organized in tandem arrays [26]. Whole genome data of diverse maize lines and wild relatives indicate that genome size variation correlates with knob content [14]. The number of knob repeats, knob size, and genomic location vary dramatically among lines. Cytological detection of knobs in recombinant inbred lines has been employed to genetically map knobs [27].

Centromeres are primarily made up with tandem satellite repeated CentC and interspersed centromeric retrotransposons of maize (CRM), both of which exhibit varying abundance across taxa [18–20]. Cytological evidence indicates that CRM elements, as the name implies, are largely located at centromeres [28]. Recently, studies using next-generation sequencing (NGS) data discovered that the abundance of CentC repeats is reduced in domesticated maize, while the contents of CRM are increased in domesticated maize, in comparison with the wild progenitor teosinte [19, 20]. Telomeres are the natural ends of eukaryotic chromosomes. Telomere repeats typically consist of 5 to 8 nucleotide highly conserved motifs, which function to recruit the proteins of the nucleoprotein complex and protect chromosomes from instability. In most plants, the conserved motif is TTTAGGG [29, 30]. Sub-telomeres are DNA sequences immediately adjacent to the telomere repeats. Hybridization, using telomere-specific probes, revealed that telomere lengths vary within a range of more than 25-fold among 22 surveyed maize inbred lines. Genetic mapping analysis mapped additional *in trans* elements that control telomere length [31]. Maize sub-telomeres consist of highly repetitive tandem sequences [32].

Here, telomere will be used as a general term for both telomere and sub-telomere repeats. Collectively, highly repetitive sequences are largely organized into clusters in maize genomes and variation in copy number is frequently observed.

NGS have provided in depth sequence data. However, accurate assessment of genome structure and dynamics of repetitive sequence evolution using large NGS datasets remains challenging due to the difficulty of unambiguous genome mapping and of accurately reconstructing repetitive sequences with high-copy number. Additionally, analysis relying on mapping reads to a reference assembly is subject to ascertainment bias. Analysis independent of a reference genome sequence could reduce biases of genome comparisons. In this study we quantify and characterize genome dissimilarity through the comparison of k-mer abundances directly determined from sequencing data. K-mers of a sequence represent all the possible subsequences of length *k*. K-mer analysis has been widely applied in many genomic analyses, such as genome assemblies, genome characterization, and metagenomic analysis [33–35]. Using B73 and Mo17 whole genome shotgun (WGS) sequencing data, we quantified the level of the difference between the two genomes at both non-repetitive and highly repetitive genome sequences. Genomic locations influencing variation in copy number at highly repetitive sequences were genetically mapped using WGS sequencing data of 280 intermated B73 and Mo17 double haploids [10]. Furthermore, highly variable k-mers in diverse lines using *Zea mays* HapMap2 WGS data [14, 15] were identified, revealing significant changes on highly repetitive sequences during maize domestication and improvement.

## Results

### K-mer analysis of genome dissimilarity between two maize inbred lines

B73 and Mo17 are two maize elite inbred lines that are widely used in maize genetic and genomic research. The two genomes have been extensively compared in both small and genome-wide scales [4–8]. However, previous studies largely relied on a reference genome, which produces systemic biases. To perform genome comparison with an unbiased k-mer method that is independent of the reference genome, two HiSeq2500 lanes of Illumina data, using PCR-free prepared DNA libraries, were generated for each of the two maize inbred lines B73 and Mo17, resulting in 450.9 and 445.3 millions of pairs of 2x125 paired-end reads, respectively. More than 99% reads were retained after the adaptor and quality trimming. The genome coverage of sequencing data (~46x) for each genome enable the employment of error correction of sequencing reads. We use abundance to represent counts of k-mer from sequencing data and use copy number to represent sequence copies in a genome. The corrected reads were subjected to 25-nt k-mer counting, resulting in approximately 749.7 and 738.7 millions of non-redundant k-mers for B73 and Mo17, respectively. The similar shapes of the distributions of k-mer abundances (Fig 1A) and the curves of cumulative contribution of k-mers with different abundances to the genomes (Fig 1B) indicate that B73 and Mo17 exhibit overall similar levels of genome complexities. The B73 and Mo17 abundance peaks are presumably located at in single-copy k-mers (<u>http://www.broadinstitute.org/software/allpaths-lg/blog/wp-content/uploads/2014/05/KmerSpectrumPrimer.pdf)</u>, which occur only once in a genome (Fig 1A). The merged B73 and Mo17 k-mer abundances form a curve with two peaks in k-mer abundances (Fig 1A). The lower abundance peak underneath the original uncombined peaks consists of k-mers specific to either B73 or Mo17, while the second higher frequency peak represents the common k-mers of the two genomes. This novel approach was employed to visualize the difference of non-repetitive genomic sequences between the two genomes. K-mer comparison indicates that only 60.9% of single-copy k-mers are shared between the two maize cultivars, leaving a remaining 39.1% of the single-copy k-mers specific to each genome (Table 1). Based on the k-mer distribution, the B73 genome size was estimated to be 2.38 Gb and consisted of 24.9% single-copy k-mers, while 2.48 Gb with 23.7% single-copy k-mers for Mo17. The B73 genome size estimation agrees with that of 2.3 Gb estimated from the B73 genome sequencing project [2]. The slightly larger estimated genome size of Mo17 versus B73 but the smaller proportion of single-copy sequences in Mo17 implies that distinct contributions of repetitive sequences to two genomes, which indeed can be observed on the curves of cumulative k-mer contribution to the genome at high abundant k-mers that are representatives of highly repetitive sequences (Fig 1B).

**Fig 1.**
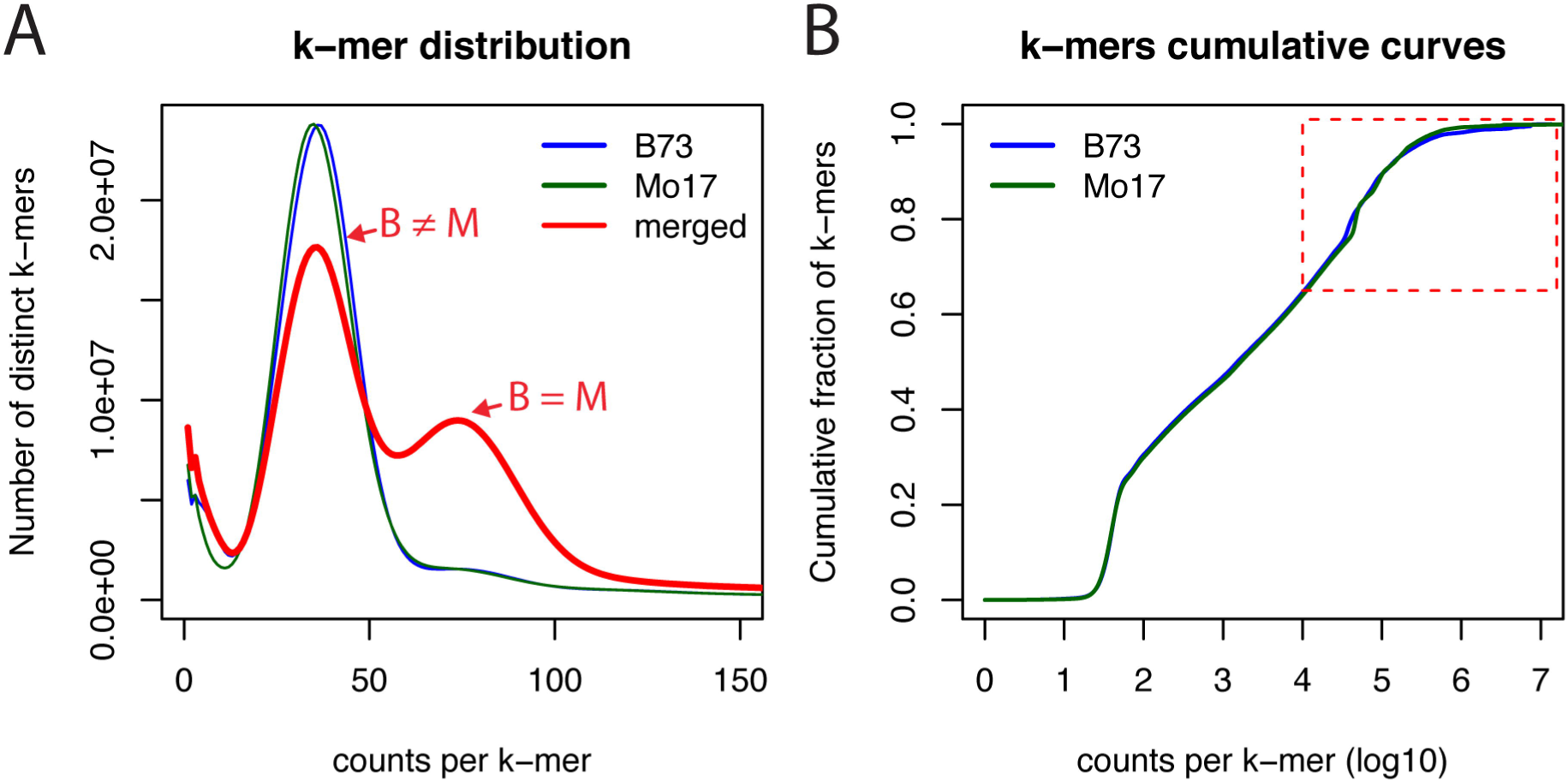
Comparison of k-mer spectra in B73 and Mo17. (A) Distributions of k-mers at different abundance in B73, Mo17, and merged B73 and Mo17. Merged k-mer counts are the total counts from both B73 and Mo17. Only the range of 1-150 on the x-axis was plotted to show the distribution of low-copy k-mers. (B) Accumulated fraction of different abundance of k-mers in each genome of B73 and Mo17. The dash-line box highlights high abundance k-mers.

**Table 1:**
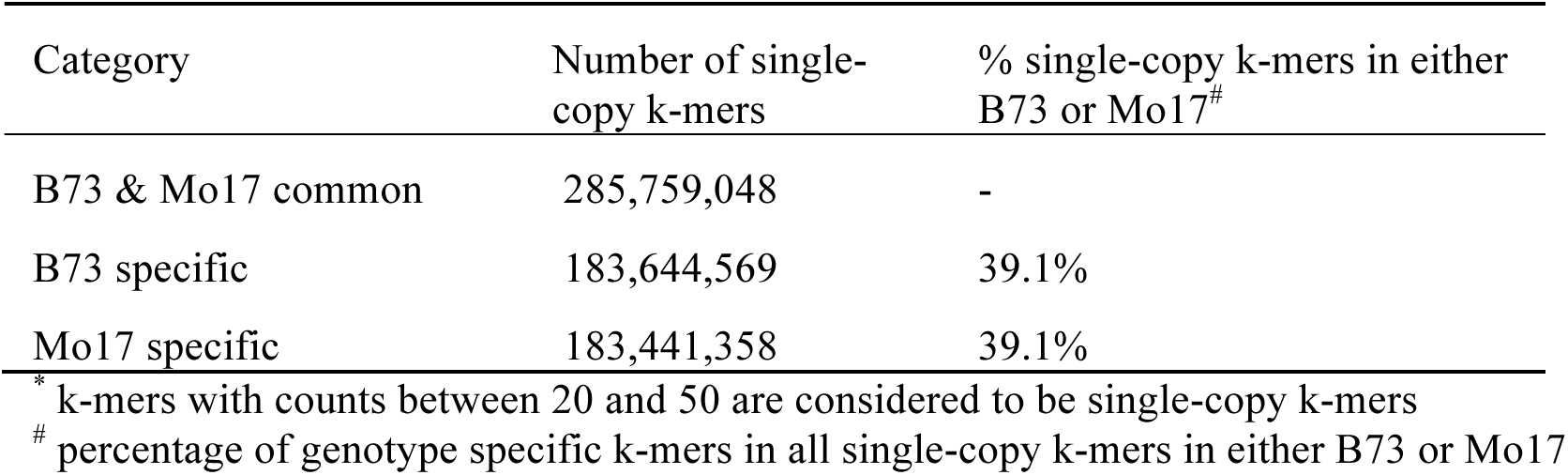
k-mers from single-copy regions in B73 and Mo17^*^

| Category | Number of single-copy k-mers | % single-copy k-mers in either B73 or Mo17<sup>#</sup> |
| --- | --- | --- |
| B73 & Mo17 common | 285,759,048 | - |
| B73 specific | 183,644,569 | 39.1% |
| Mo17 specific | 183,441,358 | 39.1% |

\* k-mers with counts between 20 and 50 are considered to be single-copy k-mers
<sup>#</sup> percentage of genotype specific k-mers in all single-copy k-mers in either B73 or Mo17

### Divergence in copy number exhibited on highly repetitive DNA sequences

Owing to the implication of the distinct constitution of high-copy genomic sequences between B73 and Mo17, highly abundant k-mers (HAKmers, N= 802,668, Table S1) in either B73 or Mo17 or both were examined. The majority of HAKmers exhibit similar abundance in the two genomes but some are highly different (Fig 2A). Functional annotation through a BLASTN of HAKmers to a *Zea mays* repeat database results in 552,371 annotated HAKmers each of which has at least one hit with the minimum e-value of 0.1. The best hit of each HAKmer was referred to as the k-mer’s functional class. The major classes include retrotransposon, knob, rDNA, CentC, telomere, and a variety of DNA transposon members (Table S2).

**Fig 2.**
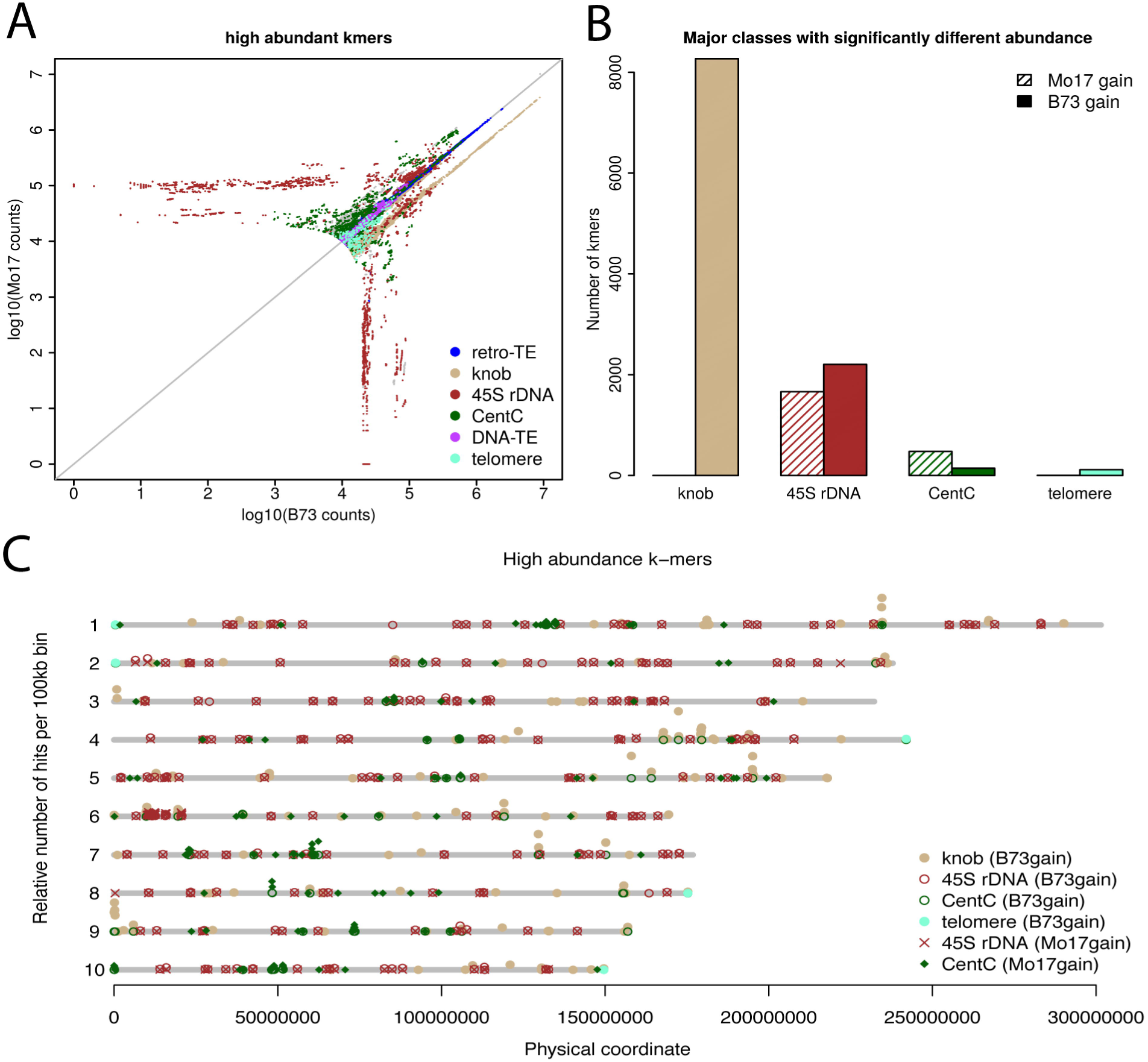
Comparison of high-copy k-mers between B73 and Mo17. (A) A scatter plot of counts of high abundant k-mers from error corrected WGS reads. K-mers were annotated by BLASTN to the maize repeat database. (B) Four major repeat classes containing k-mers that exhibit statistical significantly differential counts and at least two-fold changes between B73 and Mo17, were shown. Two types, Mo17-gain and B73-gain, respectively represent more counts in Mo17 and more counts in B73. (C) Genome-wide view of the distribution on the B73Ref3 reference genome of k-mers with differential B73 and Mo17 counts. All perfect hits each of which is end-to-end and 100% matching to the reference genome were used for determining the number of hits per bin (100 kb). The number of hits of each bin with at least 10 hits was plotted versus bin physical locations at the B73Ref3. Different functional groups were color-and shaped-coded.

*χ*^2^ statistical tests with a multiple test correction using the cutoff of 5% false discovery rate (FDR) were performed to identify HAKmers showing differential abundance between B73 and Mo17. A minimum of two-fold change in k-mer abundance was also required. As a result, 11,413 and 2,633 differential abundance HAKmers respectively showing higher abundance in B73 and Mo17 were identified, and, hereafter, referred to as B73-gain and Mo17-gain HAKmers. Four major functional annotation classes, knob, 45S rDNA, CentC, and telomere, were found in these differential abundance HAKmers (Fig 2B). Although retrotransposon derived k-mers (retrotransposon k-mers hereafter and a similar expression was applied to other classes of k-mers, e.g., 45S rDNA k-mers to represent k-mers derived from 45S rDNA) represent the largest class of HAKmers, relatively few of these differ significantly in abundance (Table S2). Many knob k-mers were identified and all belong to B73-gain k-mers, indicating more knob sequences in the B73 genome. This is consistent with the previous cytological observation that B73, but not Mo17, contains knobs at the long arms of chromosomes 5 and 7 [36, 37]. Despite the changes in the knob content detected, no differential abundance HAKmers were found to be TR-1 repeats. A similar finding was made for B73-gain telomere k-mers although the number is much smaller (Fig 2B). Moreover, a number of k-mers derived from 45S rDNA and CentC show gains in either B73 or Mo17. More 45S rDNA k-mers and less CentC k-mers showing higher abundance were identified in B73 versus Mo17. Genomic locations of these differential abundance HAKmers on the B73 genome were mapped through aligning k-mers to the B73 reference genome (B73Refv3) (Fig 2C). From the result, knob k-mers are clustered on multiple chromosomes (e.g., long arm of chromosomes 1, 4, 5, 7 and a distal short arm region at chromosome 9), CentC k-mers are largely located at or around centromeres, and telomere k-mers are identified at the ends of chromosomes 1, 2, 4, 8 and 10. 45S rDNAs k-mers are predominantly clustered at the short arm on chromosome 6, presumably the NOR. Note that such distributions based on the reference genome rely on the quality of assemblies, and the assembly quality of different regions might vary. The genome distribution plot also shows that 45S rDNA k-mers are pervasive in other genome regions in addition to the NOR (Fig 2C, Fig S1).

To understand copy numbers of different classes of highly repetitive sequences in two genomes, the total count of all the k-mers of each class was determined and normalized, which represents the relative level of repetitiveness of each class. As a result, compared to B73, approximately 55% and 22% reduction were respectively observed on knob and telomere repeats, while 71%, 34%, 25% increased on CentC, 45S rDNA and 5S rDNA, respectively, in Mo17 (Table S3). We also used abundances of the k-mers (N=3,533) from the 45S rDNA regions conserved among multiple plant species to estimate the copy number of 45S rDNA (Methods). The copy numbers of 45S rDNA in B73 and Mo17 were estimated to be around 3,658 and 5,063, respectively. Our estimation is in the range of a previous estimation of placing rRNA gene number from 2,500 to12,500 in 16 maize lines [24]. Collectively, we discovered several major classes of repetitive sequences showing differential copy number between B73 and Mo17, suggesting that two genomes experience pronounced divergence with respect to copy number of highly repetitive sequences. Because these repetitive sequences are largely clustered and tandemly arrayed, high levels of copy number variation at these loci are likely caused by insertions or deletions of large genomic segments due to aberrant crossing over or replication errors.

### Genetic mapping of genomic locations showing differential copy number of repetitive sequences between B73 and Mo17

Differential abundance of HAKmers from B73 and Mo17 results from distinct copy numbers of genomic repetitive sequences from which k-mers have originated. The segregation of such genomic sequences in a segregating population (e.g., recombinant double haploids) derived from B73 and Mo17 results in different copy number among the offspring. To map genomic locations showing the differentiation of copy number between B73 and Mo17, low-coverage WGS sequencing of 280 individuals from intermated B73xMo17 double haploids (IBM DHs) [10] was analyzed. First, the abundance of each of differential abundance HAKmers from each DH line was determined and normalized (Methods). K-mer abundance resembles a quantitative trait value, and the genomic elements contributing their genomic copy number variation can be genetically mapped using a quantitative trait locus (QTL) mapping approach (referred to as copy number variation QTL, cnvQTL, hereafter). Using a high-density genetic map developed with the same WGS data set from these 280 DH lines [10], the normalized counts of a k-mer were input as phenotypic values for a genetic mapping analysis using the R package rqtl. In total, 11,413 and 2,633 of B73-and Mo17-gain HAKmers were analyzed, respectively. To determine the cutoff of log10 likelihood ratio (LOD) of cnvQTL, each of 1,000 randomly selected HAKmers was subjected to a permutation test to determine the LOD cutoff. All of these LOD cutoffs with the 5% type I error are in between 3 and 4.

Therefore the minimum LOD of 4 was used to declare mapping cnvQTL peaks (Table S4). Only 0.3% B73-gain and 3% Mo17-gain HAKmers could not be mapped using this approach. The majority of HAKmers, 74.5% B73-gain and 83.5% Mo17-gain, were mapped to single major genomic locations, and the rest were mapped to 2-4 genomic locations.

Functional annotation analysis of these mapped HAKmers revealed distinct mapping locations for different sources of k-mers (Fig 3). For B73-gain HAKmers, knob and 45S rDNA are two major sources (Table S5). Knobs k-mers were mapped to the long arms on chromosomes 1, 5, and 7, of which the regions on chromosomes 5 and 7 were reported to have differential knobs between B73 and Mo17 [36, 37]. All 2,205 45S rDNA k-mers were mapped to around 13.5 Mb on chromosome 6 to which 11 retrotransposon k-mers were also mapped. This mapping region is located at a short arm region on chromosome 6 which exhibits a presence-and-absence variation (PAV) that was identified in previous comparative studies [7, 8]. Substantial copy gains of some type of 45S rDNA and some retrotransposons in B73 at this region indicate the long PAV segment harbors rich repetitive sequences. The differential abundance 45S rDNA k-mers are largely located at the intergenic spacer (IGS) between 18S and 26S of 45S rDNA and a small proportion are located at internal transcribed spacer (ITS) and 26S rRNA gene (Fig S2). On the same chromosome, CentC k-mers were mapped to 62.8 Mb, suggesting the two genomes contain distinct centromere compositions on chromosome 6. Moreover, telomere k-mers were mapped to the ends of short arms of chromosomes 1, 2, 4, and 5. The further analysis shows that B73 contains more copies of telomere repeats than Mo17 at chromosomes 2, 4, 5, but less copies at chromosome 1 (Table S6).

**Fig 3.**
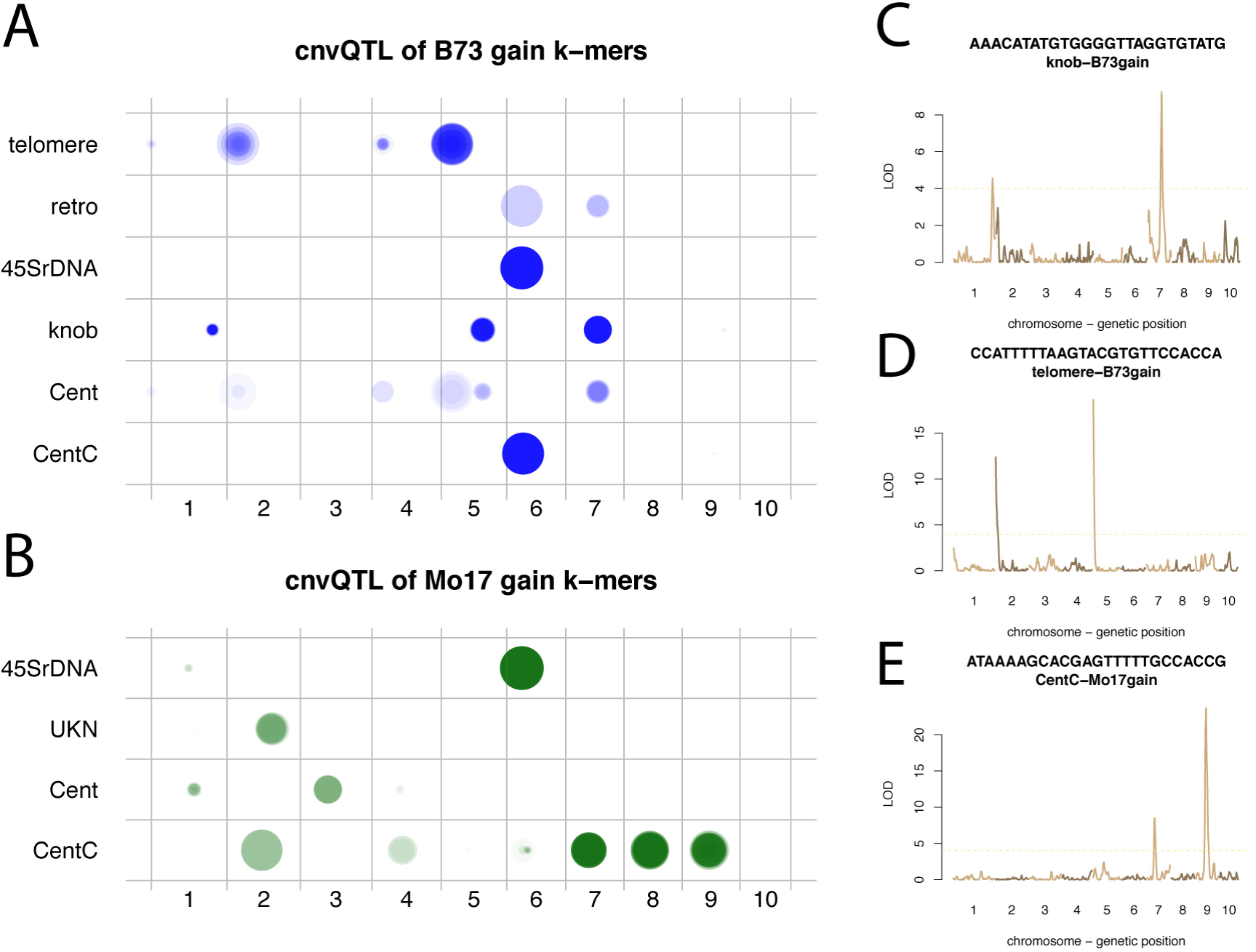
cnvQTL mapping of genomic locations contributing differential abundance of HAKmers. WGS of 280 IBM DHs was used to determine abundance of differential abundance HAKmers. A QTL approach was employed to map genomic locations influencing k-mer abundance in DH lines. (A, B) The mapping results of B73-gain HAKmers (A) and Mo17-gain HAKmers (B) were plotted for each annotated class. A mapping location of each k-mer is designated by a dot. Transparent factor (0.02) was used for a dot from each k-mer. The sizes of dots represent the logarithm 10 scaled LOD values from QTL analyses. retro, Cent, and UKN represent retrotransposon, centromere elements, and unknown elements, respectively. (C, D, E) Three examples of the QTL results of knob B73-gain (C), telomere B73-gain (D), and CentC Mo17-gain HAKmers (E), were shown.

45S rDNA and CentC are two major sources for Mo17-gain HAKmers (Table S7). Interestingly, similar to B73-gain 45S rDNA HAKmers, Mo17-gain counterparts were mapped to around 13.6 Mb on chromosome 6, although a long DNA segment on the B73 reference genome around that region is absent in Mo17. This indicates that B73 and Mo17 likely contain different versions of 45S rDNA at the NOR. Furthermore, four 5S rDNA k-mers (N=4) showing higher abundance in Mo17 were mapped to around 222.5 Mb on chromosome 2, consistent with a previous FISH result in which 5S rDNA was mapped to the distal of chromosome 2 [22]. Significantly, Mo17-gain CentC k-mers were mapped to multiple chromosomes. The centromeric regions at chromosomes 2, 4, 7, 8, and 9 contribute to varying abundance of CentC k-mers. The same k-mers can be mapped to the centromeres on multiple chromosomes, suggesting multiple centromeres co-evolved to change CentC abundance.

### Different evolutionary origins of 45S rDNAs of B73 and Mo17, likely expanded, and spread to regions other than the NOR after domestication

From differential abundance HAKmers, an extreme type of k-mer was surprisingly observed in which the k-mer was highly abundant in B73 or Mo17 but absent or very low in the other, which are referred to as genotype-specific HAKmers (Fig 4A). In total, 162 B73-specific HAKmers and 103 Mo17-specific HAKmers were obtained. These genotype-specific HAKmers were verified by using independent B73 and Mo17 WGS sequencing data [14] without error correction. Additionally, all of the B73-specific HAKmers can be perfectly aligned to the B73 reference genome, while only 3/103 Mo17 specific HAKmers were perfectly aligned to single locations at the NOR region. This result confirms, at least, that Mo17 specific HAKmers are highly abundant in Mo17 but hardly identified in the B73 genome. Interestingly, all of these genotype-specific HAKmers are annotated to the class of 45S rDNA. K-mer analysis using IBM DH lines WGS sequencing data indicates that each DH line predominated by either B73-or Mo17-specific k-mers (Fig S3). Genetic mapping analysis of both B73-and Mo17-specific HAKmers through cnvQTL shows that the NOR where 45S rDNA repeats are clustered is largely responsible for the segregation of B73-and Mo17-specific HAKmers, further suggesting that distinct types of high-copy 45S rDNAs are included at the B73 and Mo17 NORs (Fig 4B). A detailed analysis found that all these genotype-specific k-mers were mapped to the IGS of the 45S rDNA unit.

**Fig 4.**
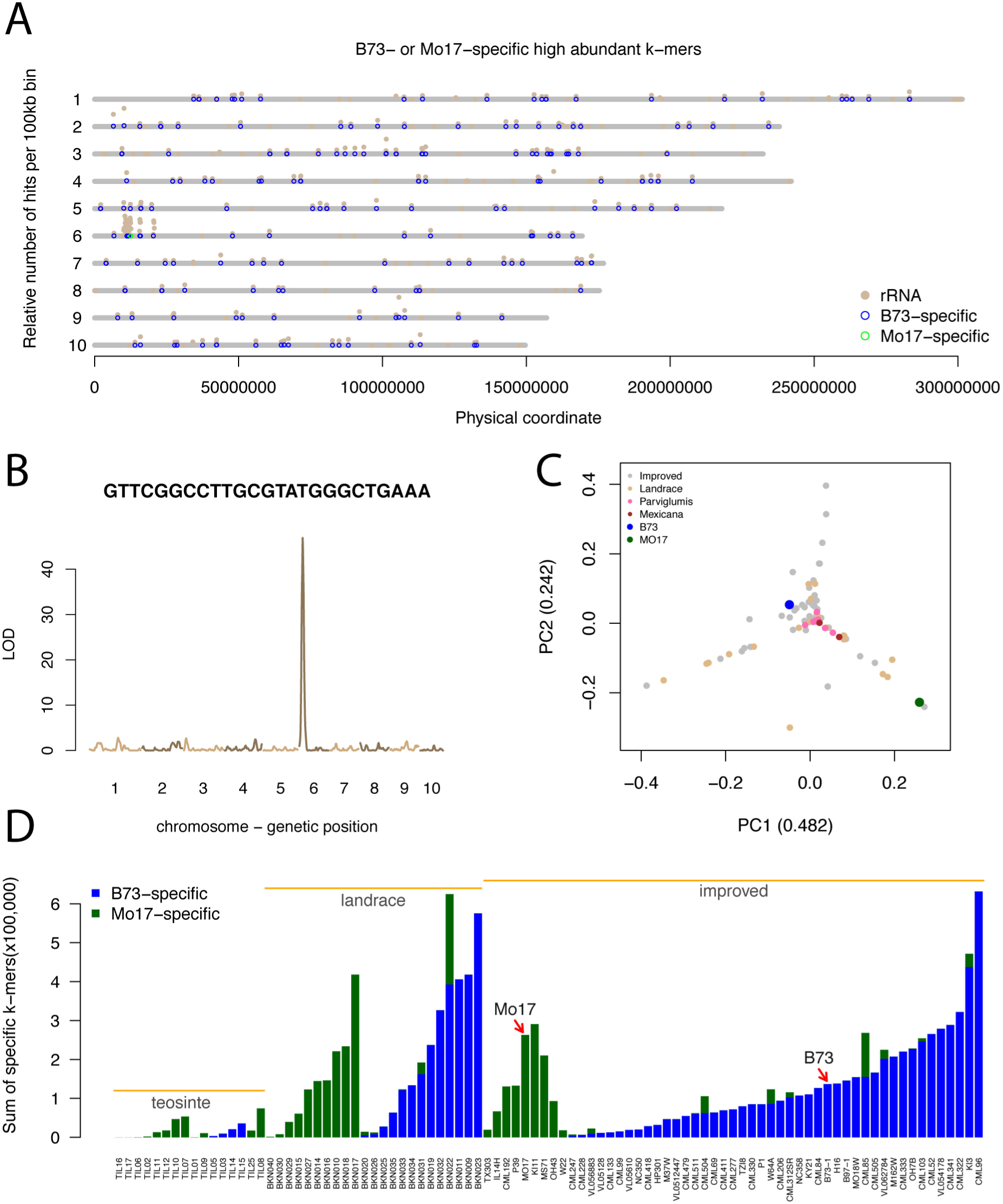
B73 or Mo17 specific HAKmers. (A) Genome-wide distributions of rDNA-related k-mers, B73-and Mo17-specific k-mers that can be perfectly aligned to the B73Ref3 reference genome. Alignment numbers per bin (100 kb) were plotted versus bin physical locations at the B73Ref3. (B) Counts of a Mo17-specific k-mer in IBM DH lines were treated as a trait and the genomic loci (or locus) contributing the levels of counts in DH lines were mapped. (C) Principal component analysis (PCA) was performed using normalized counts of each B73-or Mo17-specific k-mer in multiple teosinte, landrace, and improved maize lines. Numbers in parentheses are percentages of variation in normalized counts explained by principal component (PC) 1 and 2. (D) Sum of normalized counts of all B73-specific k-mers (blue) or Mo17-specific k-mers (green) in different lines from HapMap2 WGS sequencing data without error correction. Bars were sorted in the subspecies order, teosinte, landrace, and improved maize lines. Within each subspecies, bars were sorted by total counts of B73-specific k-mers first and then by total counts of Mo17-specific k-mers.

To understand the origin of these genotype-specific k-mers, maize HapMap2 WGS sequencing data, which includes lines from teosinte, landrace, and improved maize [14, 15], were subjected to k-mer analyses. The count of each of B73-and Mo17-specific k-mers was determined for each HapMap2 line. To account for the variation of k-mer abundance owing to non-genetic factors, such as sequencing depth and organelle DNA contamination, a novel normalization approach was developed of which normalization factors were determined by using the total counts of a set of conserved single-copy k-mers across HapMap2 lines. Briefly single-copy k-mers were first obtained from both B73 and Mo17 and the correlation of counts of each k-mer with the library sizes of all the HapMap2 lines determined. Based on the assumption that a conserved single-copy k-mer exhibits a high correlation with the sequencing library size, the top 5% k-mers (N=49,955) with highest correlation efficiencies were used to calculate the normalization factors. A principal component analysis (PCA) was performed using normalized abundances of genotype-specific HAKmers (N=265) of HapMap2 lines. At a result, the first two components (PC1 and PC2) explain 72.4% variation in normalized abundance (Fig 4C). From the PCA plot, three distinct branches were formed and teosinte lines were centralized at the intersection. Mo17 is located on the distal position of one branch but B73 is not located at any of the branches. The PCA analysis implies that not all the HapMap2 lines exhibit either of two extremely divergent patterns possessed in B73 and Mo17.

To understand the abundance of these genotype-specific HAKmers in each HapMap2 line, the total normalized counts of all the B73-and Mo17-specific HAKmers were separately determined. Total counts of the B73-and Mo17-specific HAKmers vary dramatically among the HapMap2 lines (Fig 4D). It is notable that all teosinte lines exhibit relatively low abundance, while many but not all maize lines show high abundance in total counts. This result indicates that these particular types of 45S rDNA repeats likely experienced appreciable expansion after domestication or shrinkage in teosinte and some maize lines. Evidence was also found that B73-specific k-mers are largely, but not only, located at the NOR. Indeed, the B73-specific k-mers can be identified at many locations on all the chromosomes in the B73 genome (Fig 4A).

Presumably, the scattered distribution of these k-mers across all the chromosomes is the consequence of the 45S rDNA spreading from the NOR. Moreover, all teosinte lines and the majority of maize lines contain only either B73-or Mo17-specific HAKmers, while a few landrace and improved lines consist of both. Our cnvQTL mapping result indicated that both B73-and Mo17-specific HAKmers are predominantly located at the NOR. The observed mixture of two rDNA types in some maize inbred lines are likely the consequence of heterozygous residues or recombination at the NORs, although meiotic recombination is substantially suppressed at the NOR [38]. It is also notable that the proportion of lines with B73-specific types of 45S rDNAs in the improvement levels is increased from teosinte to landrace, and from landrace to improved lines (Fig 4D), possibly due to positive selection on the NOR or nearby regions. Previous studies also suggested that this region was under selection during either domestication [15] or maize improvement [16].

### Allelic expression of 45S rDNA in hybrids of B73 and Mo17

The differences of 45S rDNA sequences in B73 and Mo17 enables the investigation of the expression of two types of 45S rDNA in the hybrid of B73 and Mo17. Messenger RNA (mRNA) is typically selected and enriched in final sequencing libraries in the regular RNA-Seq (mRNA sequencing) procedure. However, it is almost impossible to completely remove all rRNA, which allows the study of the expression of rRNA using mRNA sequencing data. Two sets of RNA-Seq data were used. One is the transcriptomic data of young maize primary roots in the B73, Mo17 and the reciprocal hybrids [39]. The other is transcriptomic data of whole kernels at 0, 3, and 5 days after pollination (DAP) and endosperms at 7, 10, and 15 DAP from reciprocal hybrids of B73 and Mo17 [40].

From both data sets, many sequences were aligned to 45S rDNA, proving that rRNA sequences remained in mRNA sequencing data. The B73-and Mo17-specific 45S rDNA k-mers can be used to trace the genotype-specific expression of 45S rDNA if their k-mer abundance could be reliably measured in RNA-Seq. However, all these genotype-specific k-mers are located at the IGS. The IGS is either not transcribed or accumulated at a level as high as the rRNA genes (5.8S, 18S, and 26S), and IGS expression therefore cannot be reliably detected. Fortunately, a single-nucleotide variant (SNV), A/T, was discovered on the 26S rRNA gene and three pairs of k-mers harboring this SNV were identified in both genomic sequencing and RNA-Seq data (Table S8). 72% and 28% B73 rDNAs carry A and T, respectively, while almost 100% of Mo17 rDNAs carry T. A-carrying rDNAs nearly completely dominated rRNA expression in primary roots of B73 (Fig 5A), suggesting that not all rDNAs, as previously reported [41], are transcribed. In Mo17, T-carrying rDNA is the only type of expressed rDNA. In the reciprocal hybrids, both types were almost identically expressed in primary roots, although in both the reciprocal hybrids the A and T types of rDNAs are unequal in abundance in their genomes (Fig 5A).

**Fig 5.**
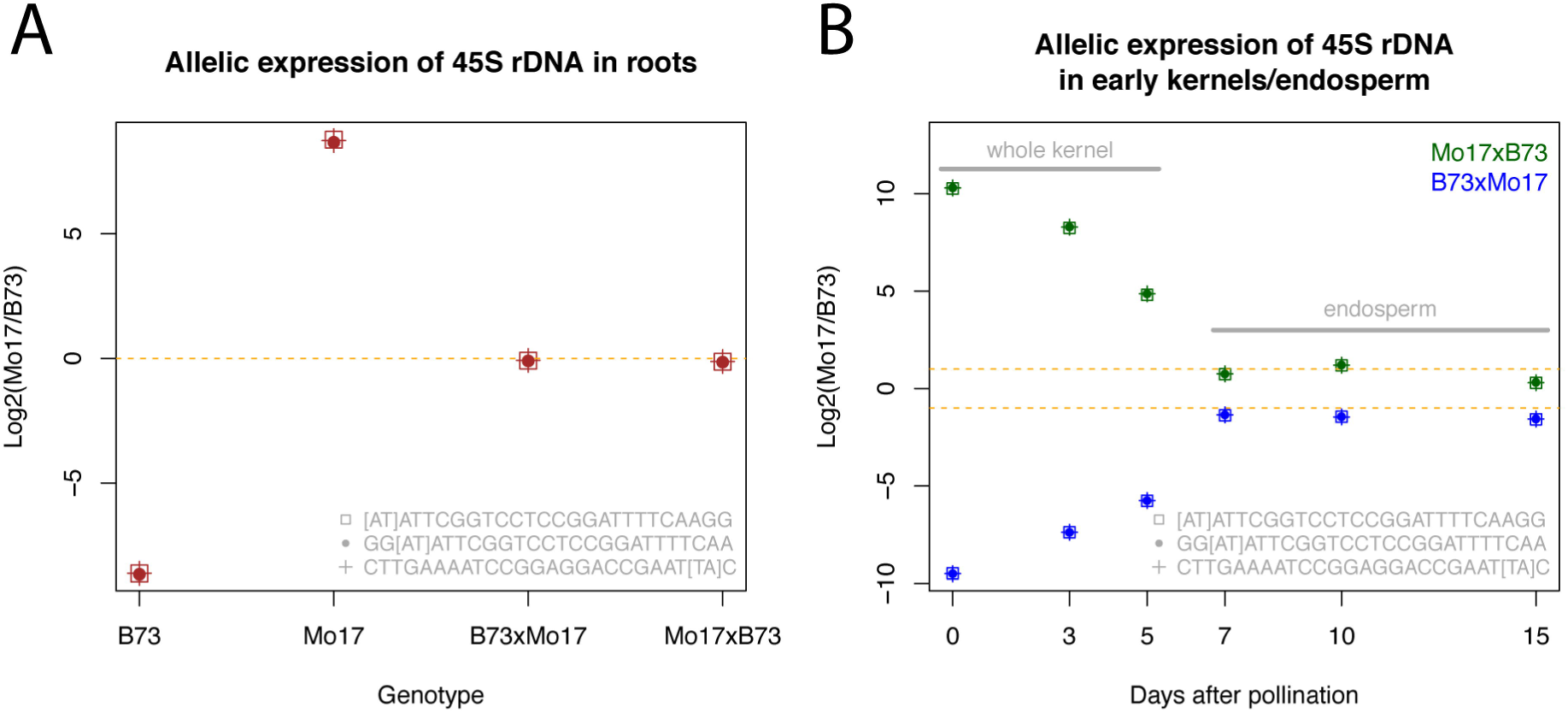
Allelic expression of 45S rDNA in hybrids of B73 and Mo17. (A) A single-nucleotide variant was identified at the 26S rRNA gene of the 45S rDNA unit. Three pairs of k-mers harboring this single-base variant were listed in the figure. Two bases within square brackets represent the allele type respectively highly enriched in B73 and Mo17 (B73 k-mer and Mo17 k-mer). The log2 of the ratio of expression abundance of each Mo17 k-mer to that of its paired B73 k-mers was plotted for four genotypes, B73, Mo17 and the reciprocal hybrids. The expression data were from maize primary root RNA-Seq. Expression abundance is the average of four biological replicates. (B) The log2 of the ratios of expression abundance of each Mo17 k-mer to that of its paired B73 k-mers were determined for samples (whole kernel or endosperm) from different developmental stages, and plotted versus the days after pollination. The expression data are from maize RNA-Seq of B73 and Mo17 reciprocal hybrids. The reciprocal hybrids were plotted in either blue (B73 as the female parent) or green (Mo17 as the female parent).

Using the time-course transcriptomic sequencing data of whole kernels and endosperms, the allelic expression levels of the SNV (A/T) on the 26S rRNA gene in the reciprocal hybrids of B73 and Mo17 were also examined (Fig 5B). As a result, detected rRNA almost entirely belong to the maternal type in whole kernels at 0 DAP. Paternal rRNA accumulation levels were gradually increased in whole kernels from 0 to 5 DAP. In endosperms at 7, 10, and 15 DAP, the ratios of maternal to paternal rRNA expression are not far from 2:1 that is the actual copy number ratio of maternal to paternal genomes, indicating that both maternal and paternal rRNA copies are expressed at equal rates in early endosperms.

**Marked changes of multiple types of highly repetitive genomic sequences during domestication and maize improvement**

The finding that B73 and Mo17 exhibit substantial variation at high-copy genomic sequences inspired an investigation of such variation among the HapMap2 lines. B73 and Mo17 are included in the HapMap2 lines but in this analysis we wanted to identify k-mers highly variable across the whole HapMap2 set, rather than the genotype-specific high abundance k-mers defined using these two inbred lines. Using the HapMap2 WGS sequencing data, k-mers showing high abundance (>1,000 counts per k-mer) in at least five HapMap2 lines but low abundance (<10 counts per k-mer) in at least five other lines were extracted, resulting in 8,462 highly variable k-mers. To examine the change of these k-mers among three evolutionary groups, teosinte, landrace, and improved, an ANOVA test was performed for each k-mer and a Bonferroni correction was conducted to account for multiple testing. As a result, 2,016 k-mers exhibit significantly differential abundance among three groups at the 5% type I error. Functional annotation through a BLASTN of k-mers to the repeat database results in 1,090 annotated k-mers (Methods). The k-mers exhibiting significantly differential abundance among evolutional groups were annotated to the functional classes of 45S rDNA, CentC, retrotransposon (copie and gypsy), and knob. The low rate (only ~54%) of k-mers that are annotated using the repeat database is because a relatively high proportion of k-mers are derived from organelle genomes, which likely reflects the diversity of organelle genomes. To focus on highly repetitive sequences from nuclear genomes, only the functionally classified k-mers were subjected to a clustering analysis using the software MCLUST [42], resulting in 12 clusters (Table S9, Fig S4). Nine major clusters were further manually grouped into two groups (Table 2, Fig 6A, B). In detail, k-mer abundance of the group 1 was significantly decreased during maize domestication and/or improvement. K-mers from this group are largely annotated as CentC (example in Fig 6C) and 45S rDNA, as well as a small number of k-mers from knob, DNA transposons, and retrotransposons (Table 2). K-mer abundance of the group 2 was substantially increased during maize domestication and/or improvement. K-mers from this group are annotated as retrotransposon members (CRM and unclassified retrotransposon) (example in Fig 6D) and 45S rDNA. The observation of 45S rDNA in both groups 1 and 2 suggests that some types of 45S rDNA sequences experienced substantial expansion while others experienced substantial shrinkage during maize domestication and improvement.

**Fig 6.**
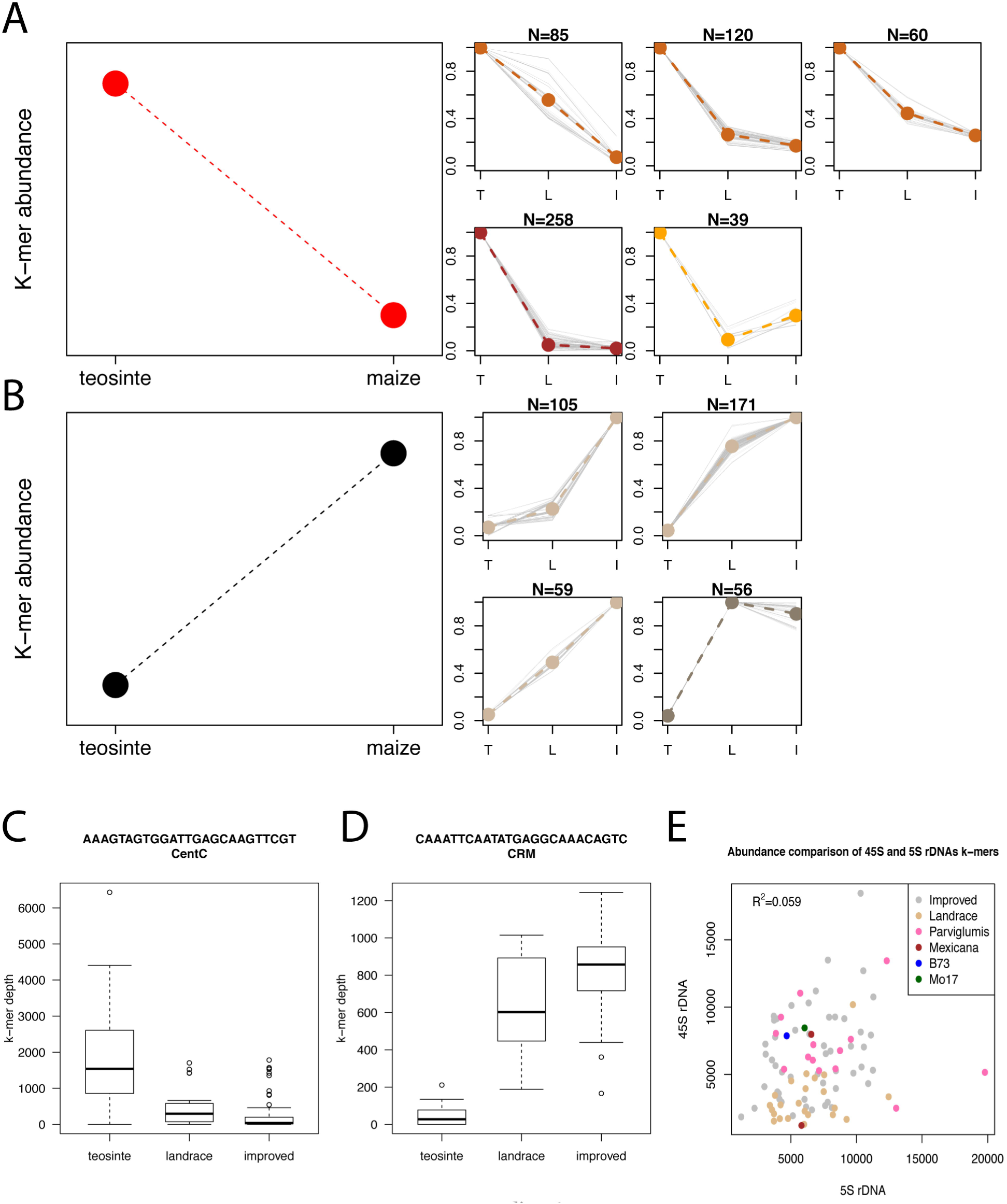
Change of k-mer abundances in teosine, landrace, and improved maize. (A, B). K-mers with significantly differential abundance in teosine, landrace, and improved maize were clustered. Nine major clusters were further manually divided into two groups. K-mers in group 1 (A) exhibit markedly higher abundance in maize relative to teosinte, while k-mers in group 2 (B) are in the opposite change. Smaller plots provide the details of the clusters in each group. Each grey line in the smaller plots represents a k-mer. Colored lines are average values from all the k-mers in each cluster. Clusters with a similar pattern were highlighted by the same color. T, L, I on the x-axes represent teosinte, landrace, and improved lines, respectively. (C, D) Boxplots of of three representative k-mers that are separately derived from CentC (C) and CRM (D). (E) The median abundance of 45S rDNA k-mers generated from the conserved 45S rDNA sequence in each HapMap2 line was plotted versus the median abundance of 5S rDNA k-mers generated from the conserved 5S rDNA sequence of the same HapMap2 line. Each dot represents a line, which is color-coded by genotype groups.

**Table 2:** Number of functionally classified k-mers in different clustering groups

| Class | Decrease during domestication | Increase during domestication |
| --- | --- | --- |
| CentC | 212 | 0 |
| CRM* | 0 | 121 |
| Knob | 12 | 0 |
| 45S rDNA | 266 | 81 |
| DNA transposon | 9 | 0 |
| Retrotransposon<sup>§</sup> | 54 | 138 |

\* k-mers were annotated unknown centromere retrotransposons
<sup>§</sup> unclassified retrotransposon

Abundance of k-mers that were generated from the conserved regions of 45S and 5S rDNA across multiple plant species was estimated for each HapMap2 line. The median of abundances of all the 45S rDNA k-mers from a HapMap2 line and the counterpart of 5S rDNA k-mers were used to represent the genomic copy number level of 45S and 5S rDNA of the line, respectively. Most landrace maize lines exhibit lower copy number than teosinte, while maize improved lines shows much higher diversity in term of 45S rDNA copy number (Fig 6E). This observation suggests there were a possible shrinkage or a strong selection on the NOR region during domestication, and a re-expansion of 45S rDNAs during improvement. No association with evolutionary groups was observed for copy number of 5S rDNAs. Additionally, the correlation of copy number of 45S and 5S rDNAs among HapMap2 lines is weak (R^2^ = 0.059), suggesting that dosage balance in genomic copy number between 45S and 5S rDNAs, which was observed in human and mouse genomes [43], was not required in *Zea* genomes.

## Discussion

This study employs a novel k-mer analysis strategy for comparative genomics. Reference-independent quantification of NGS data allows precise and unbiased comparison of the genomic constitutions, particularly highly repetitive sequences that are generally overlooked from regular analyses. Our results offer insightful information about copy number abundance, genomic locations, and evolution of highly repetitive sequences among maize genomes, and provide an unbiased genome comparative method for mining existing and incoming deluge of NGS data to gain biological insights.

### Unbiased k-mer analysis

K-mers represent all the possible subsequences of length k from a sequencing read. For genome assembly using short NGS reads, k-mers are typically generated from sequencing data to construct *de Bruijn* graphs [33]. In addition to genome assemblies, K-mer analysis has been applied to many other genomic analyses, including but not limited to characterization of repeat content and heterozygosity [34], estimation of genome size [35], evaluation of metagenomic dissimilarity [44], and identification of causal genetic variants conferring phenotypic traits [45]. Any size of k-mers can be used for k-mer analysis. Using smaller sized k-mers, sequencing data are condensed to less total k-mers, and a smaller number of k-mers are derived from single-copy regions, resulting in higher degree of information loss. Increasing size of k-mers increases both the total k-mer number and the number of single-copy k-mers, which is compromised by increased computation cost. Additionally, higher size of k-mers is more vulnerable to sequencing errors contained in sequencing reads. The impact of sequencing errors could be alleviated by error correction of sequences. The choice of k-mer length of 25 nt is an optimal size for human-sized genomes which was used in ALLPATHS-LG for analyzing k-mer abundance spectrum [46].

K-mer based methods are independent of read mapping that typically relies on a reference sequence, which allows the establishment of a fair comparison between genomes. For WGS data from either the same or different species, k-mer analysis can be directly applied to quantify the level of dissimilarity between individuals as long as WGS data are comparable. Low-coverage WGS data are sufficient to deliver reliable counts for k-mers derived from highly repetitive sequences. The critical issue is to develop a reliable normalization approach to account for non-genomic variation in data due to different sequence depths, varying levels of organelle DNAs, or contaminations from other species, particularly from microbes. In this study, we used total counts of a great number of single-copy k-mers that are conserved in the examined individuals to determine normalization factors. This normalization method is expected to well account for non-genomic variation. With high-coverage WGS data from multiple individuals, any types of genomic polymorphisms at either low or highly repetitive genomic regions would be unbiasedly represented by abundance of corresponding k-mers. In particular, copy number variation can be well captured by analyzing k-mer abundances. With that respect, one of potential applications of k-mer analysis is to perform genome-wide association with abundances of k-mers, which could retrieve some associated genetic elements that are unable to be detected using reference-based approaches. Collectively, the k-mer based approach alleviates ascertainment biases introduced by reference-based methods, and should provide the complement to many existing genome analyses.

### HAKmer copy number variation QTL mapping

Using low-coverage WGS sequencing data of the IBM DH lines, a cnvQTL genetic mapping strategy was developed to map the genomic regions determining variation of k-mer abundance among DHs. As a result, the vast majority of differential abundance HAKmers between B73 and Mo17 were confidently mapped. The success of mapping differential abundance HAKmers from a variety of sources, including 45S rDNA, CentC, knobs, and telomeres, proved the effectiveness of the cnvQTL mapping. The fact that k-mers from rDNAs, CentC, telomeres, and knobs were all mapped to the expected regions where they are physically located suggests that no recognizable *trans* elements control the segregation of these repetitive sequence copies. The lack of *trans* elements makes sense because these repetitive sequences, although they evolve rapidly, are steadily maintained in each of two maize inbred lines.

We obtained a high-resolution map identifying coordinates contributing to differences in abundance of k-mers for many types of repetitive sequences in B73 and Mo17. These mapped genomic regions accurately mark the locations of clusters of repetitive sequences and corroborate many previous findings, as well as provide additional insight into the differentiation between B73 and Mo17. For example, both B73-and Mo17-gain 45S rDNA k-mers were mapped to around 13.5 Mb (B73Ref3) on chromosome 6 where a large PAV on the order of a megabase between B73 (presence) and Mo17 (absence) has been found. The result that Mo17-gain 45S rDNA k-mers were mapped at this PAV region, presumably located on the NOR, indicates that Mo17 has distinct 45S rDNA sequences to replace the missing version of 45S rDNA at the Mo17 NOR. Moreover, some Mo17-gain 45S rDNA k-mers were mapped to 210.5 Mb at the long arm of chromosome 1 that was not discovered previously, suggesting Mo17 contains a 45S rDNA cluster with significantly elevated copy number of 45S DNA at that region. Using a set of k-mers from the 45S rDNA specific sequences that are conserved among maize, rice, and barley, we estimated that the copy number of 45S rDNA in B73 and Mo17 is 3,658 and 5,063, respectively. The Mo17-gain of 45S rDNA at chromosome 1, at least partially, explains higher copy number of 45S rDNA in Mo17 relative to B73.

Our cnvQTL mapping data genetically confirm the differential abundance of knob contents between B73 and Mo17. In addition to the long arms on chromosomes 5 and 7 that were reported previously [36, 37], a distal region (293.5 Mb) at the long arm of chromosome 1 shows higher abundance of knob repeats in B73. The reduction of knob repeats on chromosomes 1, 5, 7 primarily accounts for the 55% loss of knob repeats in Mo17. What is more, detailed differentiation in CentC and telomere sequences were revealed. The increase of CentC repeats in multiple chromosomes in Mo17 indicates a possible common driving force involved in these parallel directional changes in copy number in a genome.

### 45S rRNA expression in hybrids

Nucleolar dominance is a phenomenon specifically observed in hybrids in which the NOR of one parent are dominant over the other of which rRNA is silenced. rRNA silencing involves epigenetic modifications of chromatin [41]. To examine nucleolar dominance in hybrids, allelic expression of rRNA needs to be precisely quantified. We have showed that rRNA is well represented even in mRNA sequencing data where rRNA was selected against. The divergence of 45S rDNA sequences between B73 and Mo17 provides the possibility for examining rRNA allelic expression in their hybrids. However, most polymorphisms of 45S rDNA are located at IGS and ITS whose expression is hardly detected using the examined mRNA sequencing data. Fortunately, we identified the k-mers harboring a SNV polymorphic site on the 26S rRNA gene. The paired polymorphic k-mers are respectively, and nearly exclusively, expressed in one of B73 and Mo17 inbred lines, which sets an ideal marker to measure the expression of two types of 45S rDNA in the hybrid of B73 and Mo17. The k-mer abundance analysis indicates that Mo17 contains higher copy number of 45S rDNA than B73. Using transcriptomic sequencing data of primary roots, we observed the expression levels of rRNAs derived from two parents were equalized in both reciprocal hybrids, suggesting no nucleolar dominance occurs in the primary roots of the hybrid of B73 and Mo17 and also implying that an unknown mechanism exists to regulate dosage compensation.

Using transcriptomic sequencing data of early whole kernels and endosperms, we observed that the maternal rRNA expression is almost completely dominant in the whole kernels at 0 DAP, followed by the gradual increase of paternal rRNA expression from 0 to 5 DAP. It is not clear that inequality of maternal and paternal rRNA expression in early whole kernels is merely due to the distinct proportions of maternal and paternal genomes or its combination with the transcriptional suppression of paternal rRNA. Further examination through precise quantification of both rRNA and rDNA could address this question. Maize endosperm is a triploid, containing 2n of the material genome and 1n of the paternal genome. In early endosperms at 7, 10, and 15 DAP, the maternal rRNA expression is around twice as high as the paternal rRNA expression, indicating both maternal and paternal rRNA function, and, therefore, no nucleolar dominance was observed at the tissues examined.

### Implications for maize evolution

Maize was domesticated from a wild species teosinte (*Zea mays* ssp. *parviglumis*) approximately 9-10 thousands years ago [12, 47]. Genetic evidence supports a single domestication and the post-domestication introgression from other wild relatives including *Zea mays* ssp. *Mexicana* [12, 15, 48]. The two distinct versions of 45S rDNA repeats traced by B73-and Mo17-specific k-mers at the NOR can be identified in different teosinte lines, indicating maize NORs originated from multiple ancient sources. The lower abundance of B73-and Mo17-specific k-mers in all examined teosinte but higher abundance in most landraces and improved maize lines suggests an expansion of certain types of rDNA repeats after domestication. Our observation that identical genotype-specific sequences are spread throughout the entire genome also raises interesting questions about the evolutionary past and origin of these sequences in relation to the NOR. Given evidence for a single domestication event and our observation of the local expansion of genotype-specific 45S rDNA sequences during maize domestication and improvement, flow of rDNA repeats away from the NOR following domestication is a more likely hypothesis. While the translocating mechanism can be either RNA-or DNA-mediated, our observation that spread regions consist of tandem arrays of intact 45S rDNA repeats suggests that this translocating mechanism is likely DNA-mediated. Spreading phenomena were observed for knob repeats and centromere retrotransposon members in both our results (Fig. 2) and previous studies [18, 27, 49, 50]. Spreading sequences might serve as seeds that could eventually form new clusters of repetitive sequences, such as nascent knobs or NORs.

To further characterize flux of repetitive DNA during evolution, we identified k-mer sequences showing strikingly differential abundance among three groups, teosinte, landraces, and improved lines. Nearly all of these differential abundance k-mers displayed distinct patterns of either increase or decrease in abundance from teosinte to maize. rDNA k-mers make up the largest class of differential abundance k-mers. While 83 45S rDNA k-mers showed increasing abundance during this evolutionary time-frame, 266 showed marked loss. Additional analysis of relative copy numbers of 45S rDNA of HapMap2 lines also showed shrinkages and expansions of 45S rDNA repeats from teosinte to maize lines. In contrast, all differential abundance CentC k-mers were observed to decrease in abundance, strongly suggesting the shrinkage of CentC during domestication. The reverse trend is seen for CRM k-mers, which are dramatically elevated during domestication. This result replicates similar findings discussed in two recent centromere publications [19, 20]. In addition, other retrotransposon members vary greatly among historical groups. Increasing evidence shows that transposons play important roles in adaptation and evolution [51, 52]. The dramatic change in copy number of transposon elements during maize domestication could affect transcription and gene function by disrupting genes via direct integration in functional genic regions, providing new regulatory elements, and spreading epigenetic status to nearby genes [53, 54]. In summary, our k-mer analyses offers a single-base resolution to trace dynamics of *Zea mays* genomes which has been appreciated through cytogenetics, molecular, genetics, and genomics studies, providing valuable insights into the contents and organization of highly repetitive sequences in maize.

## Methods

### Plant materials and extraction of nucleus genomic DNA

Two sources of B73 (PI 550473) were used, including seeds from Patrick Schnable laboratory and North Central Regional Plant Introduction Station (NCRPIS). All Mo17 (PI 558532) seeds were originated from NCRPIS. Seeds of two genotypes were geminated and grown in growth chamber at 28°C, with a photoperiod of 14:10h (light:dark). 15~20 grams of fresh leaves of seedlings at 2-3 leaf-stage were harvested, frozen in liquid nitrogen, and homogenized with liquid nitrogen to fine powder. The nuclei were isolated using a protocol modified from Zhang’s approach [55], followed by using the Qiagen DNeasy Plant Mini Kit protocol to extract nucleus DNA.

### WGS sequencing of B73 and Mo17

Genomic DNAs from nuclei were used for PCR-free library preparation. Two replicates of each of B73 and Mo17 were whole genome shotgun sequenced with one sample per lane in HiSeq2000. 2x125 bp paired-end data were generated. Sequencing was conducted at BGI Genomics Co., Ltd., Shenzhen, China.

### Error correction and genome size estimation

B73 and Mo17 whole genome sequences were trimmed to remove adaptor contaminations and low quality sequences with Trimmomatic (version 3.2) [56]. The clean data were subjected to error correction using the error correction module (ErrorCorrectReads.pl) in ALLPATHS-LG [46] with the parameters of “PHRED_ENCODING=33 PLOIDY=1”. Genome size was estimated during the procedure of error correction.

### K-mer counting

Corrected sequences were subjected to k-mer counting using the count function in JELLYFISH [57] with the k-mer size of 25 nt.

### Estimation of genomic copy number of 45S rDNAs in B73 and Mo17

Quantification of rDNA copy number was performed using k-mers generated from the conserved regions of 45S rDNA among maize, rice, and barley. K-mers were aligned to the *Zea mays* repeat database (TIGR_Zea_Repeats.v3.0) to exclude any k-mers aligning to non-45S rDNA repeats, and to the B73Ref3 mitochondrial and plastid sequences to exclude k-mers that are not exclusively nuclear. Abundance of the 45S rDNA k-mers was evaluated for each B73 and Mo17. Abundances of these conserved k-mers in B73 and Mo17 were normalized by division by the respective estimated abundances for single-copy k-mers in order to estimate the number of 45S rDNA repeats in each genome. The median value of all conserved k-mers was the estimation of the rDNA copy number.

### Identification of HAKmers with significant abundance between B73 and Mo17

High-abundance k-mers (HAKmers) in B73 or Mo17 were extracted, each of which is required to have at least 20,000 of total of B73 and Mo17 counts. A *χ*^2^ statistical test for each HAKmer was performed to test the null hypothesis of no relationship between k-mer counts and the genotypes (B73 and Mo17). P-values of all HAKmers were corrected to account for multiple tests [58]. The differential abundance of HAKmers were declared if adjusted p-values are smaller than 5% and fold change in k-mer abundance between B73 and Mo17 is not less than 2.

### Functional annotation of HAKmers

The *Zea mays* repeat database (TIGR_Zea_Repeats.v3.0) was downloaded from the plant repeat database that is currently maintained by Michigan State University (plantrepeats.plantbiology.msu.edu). BLASTN was performed with the word size of 12 to identify hits in the *Zea mays* repeat database for each HAKmer. The top hit with the e-value cutoff of 0.1 was referred to as the functional annotation.

### K-mer mapping to the B73 reference genome

K-mer mapping to the B73 reference genome (B73Ref3) was conducted by using Bowtie (version 1.1.2) to identify all possible perfect hits.

### Genetic mapping of HAKmers via cnvQTL

Resequencing data of 280 DH lines of the IBM Syn10 population used to build an ultra-high density genetic map [10] were trimmed with Trimmomatic (version 3.2) [56].

Remaining clean reads were subjected to k-mer counting with JELLYFISH. The k-mer size is 25 nt. The abundance of each HAKmer with differential abundance in B73 and Mo17 was determined in each DH line. The total counts (*C*) of a million of randomly selected B73 and Mo17 common single-copy k-mers in each DH line were determined. The normalization factor for the *i*th line was calculated by using the formula *NC_i_*/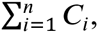 where *N* is the total number of IBM DH lines. The designation single-copy was determined by k-mer abundance from whole genome sequencing data for both B73 and Mo17 and confirmed by alignments to the B73ref3. Normalized abundance of a HAKmer was treated as a quantitative trait. For each HAKmer, a genetic mapping resembling a QTL detection implemented in an R package rqtl [59] was performed to identify the genomic locations contributing the HAKmer abundance.

### Identification of B73-and Mo17-specific HAKmers

To identify extremely unbalanced HAKmers that show extremely low abundance in one of two datasets from B73 and Mo17, the maximum number of 10 was used as the cutoff. Note that the minimum total abundance from B73 and Mo17 is 20,000 for HAKmers. If a HAKmer exhibits extremely low abundance (<=10) in one genotype, it must be high (>19,990) in the other genotype. An extremely unbalanced HAKmer of which only one genotype, B73 or Mo17, showing high abundance is called B73 or Mo17 specific HAKmers.

### HapMap2 data and k-mer analysis

Resequencing data of *Zea mays* HapMap2 lines [14, 15] were downloaded and trimmed with Trimmomatic (version 3.2), followed by 25 nt k-mer analysis using JELLYFISH. To make comparable k-mer abundances in different lines, a novel normalization method was developed. In this method, a set of “conserved single-copy k-mers” across HapMap2 lines was identified, which are single-copy in almost all lines. For each of these k-mers, k-mer abundances of HapMap2 lines should show a high correlation with their sequencing library sizes. In detail, the k-mer abundance of each HapMap2 line was determined for each of one million of B73 and Mo17 common single-copy k-mers that we identified. For each k-mer, a correlation of k-mer abundances of HapMap2 lines with their library sizes was calculated. The top 5% k-mers with the highest correlations (N=49,955) were selected, which are deemed as “conserved single-copy k-mers”. The total counts (*C*) of conserved single-copy k-mers per Hapmap2 line were determined. The normalization factor for the *i*th line was calculated by using the formula 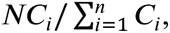 where *N* is the total number of HapMap2 lines.

### PCA of k-mer abundance of B73 and Mo17 specific k-mers in HapMap2

PCA was performed using normalized k-mer abundances of B73 and Mo17 specific k-mers in HapMap2. The R function of *prcomp* was used for the PCA.

### Allelic expression of rRNA in hybrids of B73 and Mo17

The RNA-Seq data of young maize primary roots in the B73, Mo17 and their reciprocal hybrids [39] and the time-course sequencing RNA-Seq data of whole kernels at 0, 3, and 5 DAP and endosperms at 7, 10, and 15 DAP from reciprocal hybrids of B73 and Mo17 [40] were downloaded. Sequencing reads were subjected to quality, adaptor trimming, and k-mer counting with the size of 25 nt. The expression abundance of 45S rDNA k-mers harboring a polymorphic site was used to assess allelic expression.

### Identification of highly variable k-mers in *Zea mays*

Abundances of k-mers were determined in each HapMap2 line. K-mer abundances were normalized using normalization factors calculated from a “conserved single-copy k-mers”. Highly variable k-mers were extracted using the hard-filtering criteria that require >1,000 counts per k-mer per line in at least five HapMap2 lines but <10 counts in at least five other lines.

### Identification of highly variable k-mers with significant differential abundance among evolutionary groups

Normalized counts of each k-mer for all HapMap2 lines were subjected to an ANOVA test. The genotype variable has three levels: teosinte, landrace, and improved. The null hypothesis is that k-mer abundances are independent of the genotype evolutionary groups. Then the Bonferroni approach was applied for multiple test correction at the 5% type I error.

### MCLUST to classify highly variable k-mers showing significantly differential abundance among evolutionary groups

K-mers exhibiting significant differential abundance among three genotype groups were subjected to a clustering analysis using MCLUST [42]. For each k-mer, each count was scaled by being divided by the maximum count value of this k-mer. Scaled counts of k-mers were then used for the clustering using the parameters of “G=1:12, modelNames ='EEE'”.

### Estimation of relative genomic copy number of rDNAs in HapMap2 lines

The 45S rDNA k-mers used to estimate 45S rDNA copy number in B73 and Mo17 were used to estimate relative copy number level of each HapMap2 line. In each line, the median abundance value of the k-mers represents the 45S rDNA copy number. The same method was used to determine 5S rDNA copy number level. The 5S rDNA k-mers were derived from the 5S rDNA sequence that is conserved among maize, rice, and wheat and were not aligned to B73 organelle genomes and other repetitive sequences.

## Data access

B73 and Mo17 Illumina sequencing data have been deposited at Sequence Read Archive (SRA accession number: SRP082260).

## Acknowledgments

We thank Drs. Nathan Springer, Eduard Akhunov, and Jeffrey Ross-Ibarra for discussions and valuable suggestions. We thank the support from the Kansas Agricultural Experiment Station of Kansas State University. This is contribution number 17-072-J from the Kansas Agricultural Experiment Station. We thank the support from the Agricultural Science and Technology Innovation Program of CAAS.

